# Sequence elimination in hybrid offspring of wheat-*Agropyron cristatum* (L.) Gaertn introgression line Pubing3504 × common wheat cultivar Jing4839

**DOI:** 10.1101/067504

**Authors:** Wu Xiaoyang, Chen Dan, Lu Yuqing, Zhang Jinpeng, Liu Weihua, Yang Xinming, Li Xiuquan, Du Juan, Li Lihui

**Author notes:** Corresponding author: Li Lihui.

## Abstract

Sequence elimination is one of main reasons for homologous chromosome differentiation in common wheat. Sequence elimination can occur in genome-specific sequences, chromosome-specific sequences, and repeat sequences in the wheat genome. Genetic polymorphism loci in chromosome-specific sequences can be used to develop molecular markers including simple sequence repeats (SSRs), insertions and deletions, and single nucleotide polymorphisms (SNPs). Pubing3504 is a *wheat-Agropyron cristatum* (L.) Gaertn introgression line, and Jing4839 is a common wheat cultivar. Assessment of their recombinant inbred line (RIL) population using 120 pairs of SSR markers covering all wheat chromosomes indicated that sequence elimination occurred at the short arm of chromosome 1A (1AS). We developed 13 pairs of new co-dominant SSR markers and constructed a genetic linkage map of 1AS; we found that the segment with sequence elimination is from *SSR110* to the end of 1AS. We further developed 10 pairs of dominant SNP markers of Pubing3504, 10 pairs of dominant SNP markers of Jing4839, and 10 pairs of primers designed in SNP flanking sequences to assess RILs. We found that all chromosome segments with sequence elimination came from Jing4839. The sequence elimination occurred in SSR loci, SNP loci, and coding sequences. There was no homologous recombination in the chromosome segment with sequence elimination. We suggest that sequence elimination causes the differentiation of chromosomes and the chromosome differentiation affects the homologous pairing at the chromosome segment in meiosis, which further affects the occurrence of homologous recombination at the chromosome segment.

## INTRODUCTION

Allopolyploidization is one of the main forces driving the evolution of plant genomes, inducing structure–function and modification differentiation in plant genomes (Wendel 2000; Feldman and Levy 2005; Otto 2007; Leitch and Leitch 2008). This differentiation is known as “genome shock” (McClintock 1993). “Genome shock” widely exists in natural and artificial interspecific hybridization. Allopolyploidization brings rapid changes in the genome of hybrid offspring (Comai 2000; Chen 2007). These genome changes include chromosomal rearrangements, chromatin remodeling, gene expression alteration, DNA methylation, histone modification, and reactivation of transposable elements (Kashkush et al. 2003; Pontes et al. 2003; Pontes et al. 2004; Adams and Wendel 2005; Gaeta et al. 2007; Ha et al. 2009).

Common wheat (*Triticum aestivum*) is an allohexaploid with genome composition AABBDD, which has experienced at least two interspecific hybridizations in the process of its evolution. The genetic polymorphism of the wheat genome not only comes from the accumulation of random mutations, but the changes also largely come from allopolyploidization (Feldman and Levy 2005; Feldman and Levy 2012). During the evolution of common wheat, the low-copy sequences (Liu et al. 1998a) and the repeated sequences (Salina et al. 2004; Han et al. 2005) in the genome have been eliminated in large scale. Sequence elimination is not a random event, and the elimination of coding and non-coding sequences results in the generation of the chromosome differentiation and new genetic diversity in the wheat genome. Furthermore, these eliminated sequences are within the recognition site of homologous chromosome pairing in meiosis (Feldman et al. 1997). In the process of allopolyploidization, sequence elimination is usually accompanied by wide changes in epigenetic modification and gene expression (Shaked et al. 2001; Kashkush et al. 2002; Kashkush et al. 2003; Levy and Feldman 2004; Feldman and Levy 2005; Feldman and Levy 2009).

Genomic and transcriptional changes after allopolyploidization exhibit genome preference (Ozkan et al. 2001; Ma et al. 2004). In hybrid progeny between *Aegilops sharonensis* (S^l^S^l^) and *Aegilops umbellulata* (S^u^S^u^), 14% of the *Ae. sharonensis* genome sequence was eliminated, but only 0.5% of the *Ae. umbellulata* genome sequence was eliminated. In hybrid progeny between *Ae. sharonensis* and *Triticum monococcum*, the eliminated sequence in the *T. monococcum* genome was twice as large as that of *Ae. Sharonensis* (Shaked et al. 2001). Analysis of genome-wide transcription of the hybridization progeny and the parents revealed that the patterns of gene expression in synthetic hexaploid wheat are homoeolog specific, non-additive, and parentally dominant (Akhunov et al. 2010; Chagué et al. 2010; Qi et al. 2012). In allohexaploid wheat, morphology and environmental adaptability are usually controlled by only one genome among the A, B, and D genomes (Peng et al. 2003; Nalam et al. 2006; Feldman and Levy 2012). However, the mechanism of its establishment and maintenance is still not clear.

*Agropyron cristatum* L. Gaertn is one of the main plant species of *Triticeae*. Its chromosome composition is seven as the base number, and the chromosomes are diploid and tetraploid (Dong 1993). Our laboratory successfully completed the hybridization between tetraploid *A. cristatum* and common wheat, and subsequently, a large number of introgression lines were created (Wang et al. 2011; Dan et al. 2012; Lu et al. 2015). The chromosome number of these materials is consistent with common wheat, but their alien genetic component cannot be effectively detected by cytology or the molecular marker method. For this reason, we know little about the changes in the genomes of these introgression lines. However, it is clear that these introgression lines have undergone more allopolyploidization than has common wheat. In addition, many of these introgression lines show excellent breeding traits including multiple grains and extensive resistance to wheat diseases and insect pests. Some materials have been used in breeding and gene mapping.

Pubing3504 is one of the *wheat-Agropyron cristatum* (L.) Gaertn introgression lines, and a recombinant inbred line (RIL) population was established by the hybridization of Jing4839 (a common wheat) and Pubing3504. The sequence elimination at the short arm of chromosome 1A (1AS) was found from the RIL population. In this study, we developed a series of molecular markers for the elimination of chromosome segments used chromosome-specific sequences to determine the range and the origin of the chromosome segment with sequence elimination. In addition, we further studied the effect of sequence elimination on the coding sequence and homologous recombination.

## MATERIALS AND METHODS

### Plant materials

The *wheat-Agropyron cristatum* (L.) Gaertn introgression line Pubing3504, common wheat cultivar Jing4839, and their RIL population (F8) including 336 lines were used for this study.

### Development of molecular markers

We developed new simple sequence repeat (SSR) markers in 1AS. The diploid wild einkorn wheat (*Triticum urartu*) is the progenitor species of the A genome. Its scaffold sequence bin mapping in 1AS was used in this study to develop SSR markers (Ling et al. 2013). A high-density genetic linkage map was constructed using the wheat 90K chip (Wang et al. 2014). We further obtained a scaffold sequence using the chip-probe sequence in the 1AS blast scaffold sequence of *T. urartu*. The screening standard was identities higher than 95%. The Perl script of S. Cartinhour (Temnykh et al. 2000) was used to search for SSRs in the scaffold sequence, and SSR sites with repeats of >10 were used to design SSR markers.

We developed new single nucleotide polymorphism (SNP) markers in 1AS using the allele-specific PCR (AS-PCR) technique (Wangkumhang et al. 2007). Two methods were used to obtain SNP sites between the parents; the first method was wheat 90K chip detection, and the second was RNA sequencing described in *transcriptome analysis*. The chromosomal locus information of the chip-SNP was derived from the reference genetic map of the 90K chip. The flanking sequence information of the chip-SNP was obtained using the probe sequence of the 90K chip blast scaffold sequence of *T. urartu*. The screening standard was identity values higher than 95%. The chromosomal loci and sequence information of RNA-sequencing SNP were derived from transcriptome analysis. Primer3 was used for primer design (Koressaar and Remm 2007).

Whole-genome sequencing was completed for China Spring (CS), a common wheat cultivar (Mayer et al. 2014). The cDNA sequence, scaffold sequence, and ensemble sequence (http://plants.ensembl.org/Triticum_aestivum/Info/Index) of CS were used in this study. The positions of markers in ensemble chromosome 1A were obtained by marker sequence as query to blast ensemble chromosome 1A sequence.

### Detection of molecular markers and construction of the genetic map

Genomic DNA was extracted from young leaves of all plant materials (Allen et al. 2006). We used 120 polymorphic SSR markers published in GrainGenes 2.0 (http://wheat.pw.usda.gov/GG2/index.shtml) to genotype the RIL population. PCR amplification was performed in reactions (10 μL final volume) containing 60 ng template DNA, 1 U Taq polymerase, 1× PCR buffer, 200 μmol/L dNTP, and 0.25 μmol/L of primers. DNA was amplified for 5 min at 94°, then 35 cycles of 60 s at 94°, 60 s at 50–65°, and 60 s at 72°; and 10 min at 72° for a final extension. The PCR products were separated on 8% polyacrylamide denaturing gels (Acr:Bis = 19:1) and visualized with silver staining. All marker data were scored by visual inspection, and ambiguous bands were not scored. The genetic map was constructed using QTL IciMapping software (Meng et al. 2015).

### Transcriptome analysis

The scaffold sequence of 1AS was used as the reference in this study. Young leaves of two parents and a sequence elimination line from the RILs were used for transcriptome sequencing. Reads of the two parents were mapped to the reference sequence using bwa (Li and Durbin 2009). SAMtools and BCFtools were used for SNP calling (Li et al. 2009; Li 2011). We further obtained SNPs between Pubing3504 and Jing4839 through the comparison of the parental SNP calling results. Flanking sequences (60 bp) of these SNPs were used to blat (Kent 2002) reads of the sequence elimination line. The SNPs that did not appear in reads of the sequence elimination line were used to develop SNP markers.

## RESULTS

### Detection of sequence elimination using co-dominant SSR markers

To map the breeding traits of Pubing3504, we assessed 336 RIL lines constructed from the hybrid combination between Pubing3504 and Jing4839. We obtained 120 pairs of polymorphic markers covering the 21 pairs of wheat chromosomes using parental genomic DNA to screen SSR markers published at the GrainGenes website. All 120 pairs of polymorphic markers were used to genotype the RIL population. In the RIL population, the normal segregation ratio of the parental genotypes is 1:1. In the genotype result, however, some markers exhibited abnormal separation; the number of lines that had the Pubing3504 genotype was significantly higher than that of lines that had the Jing4839 genotype, and the number of lines that had missing genotype was higher than normal. These abnormally separated markers were *Xwmc24, Xpsp2999, Xbarc263, Xcwm75*, and *Xcfa2153*, all located in 1AS (Table 1).

**Table 1.** The detection result of the RIL population using 120 pairs of SSR markers in GrainGenes 2.0.

| Marker name | Chromosome | Number of Pubing3504<br>genotypes | Number of J4839<br>genotypes | Number of missing<br>genotypes | $\chi^2$ value |
| --- | --- | --- | --- | --- | --- |
| *Xwmc24* | 1A | 227 | 33 | 76 | 144.94 |
| *Xbarc263* | 1A | 223 | 37 | 76 | 131.22 |
| *Xpsp2999* | 1A | 225 | 35 | 76 | 136.99 |
| *Xcwm75* | 1A | 224 | 36 | 76 | 132.14 |
| *Xcfa2153* | 1A | 226 | 34 | 76 | 141.78 |
| *Xbarc105* | 7D | 176 | 159 | 1 | 0.76 |
| *Xbarc117* | 5A | 175 | 161 | 0 | 0.58 |
| *Xbarc12* | 3A | 161 | 175 | 0 | 0.58 |
| *Xbarc121* | 7A\7D | 171 | 163 | 2 | 0.30 |
| *Xbarc142* | 5B | 184 | 152 | 0 | 3.05 |
| *Xbarc147* | 3B | 177 | 159 | 0 | 0.96 |
| *Xbarc176* | 7B | 184 | 151 | 1 | 3.44 |
| *Xbarc177* | 5D | 171 | 165 | 0 | 0.11 |
| *Xbarc186* | 5A | 167 | 169 | 0 | 0.01 |
| *Xbarc21* | 6D | 189 | 147 | 0 | 5.25 |
| *Xbarc232* | 5B | 175 | 159 | 2 | 0.96 |
| *Xbarc252* | 7D | 185 | 151 | 0 | 3.44 |
| *Xbarc292* | 7A | 175 | 161 | 0 | 0.58 |
| *Xbarc310* | 3A | 172 | 164 | 0 | 0.19 |
| *Xbarc322* | 5D | 168 | 168 | 0 | 0.00 |
| *Xbarc3220* | NA | 164 | 170 | 2 | 0.19 |
| *Xbarc324* | 3A | 174 | 162 | 0 | 0.43 |
| *Xbarc49* | 7A | 181 | 155 | 0 | 2.01 |
| *Xbarc50* | 7B | 180 | 156 | 0 | 1.71 |
| *Xbarc59* | 5B | 171 | 162 | 3 | 0.43 |
| *Xbarc67* | 3A | 180 | 156 | 0 | 1.71 |
| *Xbarc68* | 6B | 175 | 161 | 0 | 0.58 |
| *Xbarc70* | 7A | 190 | 146 | 0 | 5.76 |
| *Xbarc72* | 7B | 175 | 161 | 0 | 0.58 |
| *Xbarc75* | 3B | 166 | 170 | 0 | 0.05 |
| *Xbarc90* | 7B | 171 | 165 | 0 | 0.11 |
| *Xcfa161* | NA | 172 | 162 | 2 | 0.43 |
| *Xcfa2028* | 7A\7B | 162 | 174 | 0 | 0.43 |
| *Xcfa2147* | 1A\1B\1D | 169 | 167 | 0 | 0.01 |
| *Xcfa2219* | 1A | 170 | 166 | 0 | 0.05 |
| *Xcfa2256* | 4A | 180 | 151 | 5 | 3.44 |
| *Xcfa2263* | 2A | 157 | 179 | 0 | 1.44 |
| *Xcfd49* | 6D | 161 | 174 | 1 | 0.58 |
| *Xcfd54* | 4B\4D | 173 | 163 | 0 | 0.30 |
| *Xcfd55* | 3D | 168 | 165 | 3 | 0.00 |
| *Xcfd8* | 5D | 166 | 170 | 0 | 0.05 |
| *Xcfd84* | 4D | 172 | 164 | 0 | 0.19 |
| *Xgwm122* | 2A | 165 | 169 | 2 | 0.11 |
| *Xgwm126* | 5A | 182 | 154 | 0 | 2.33 |
| *Xgwm132* | 6B\6D | 177 | 159 | 0 | 0.96 |
| *Xgwm140* | 1B | 173 | 163 | 0 | 0.30 |
| *Xgwm149* | 4B\4D | 162 | 174 | 0 | 0.43 |
| *Xgwm155* | 3A | 163 | 173 | 0 | 0.30 |
| *Xgwm182* | 5D | 179 | 157 | 0 | 1.44 |
| *Xgwm213* | 5B | 183 | 153 | 0 | 2.68 |
| *Xgwm219* | 6B | 148 | 188 | 0 | 4.76 |
| *Xgwm232* | 1D | 157 | 179 | 0 | 1.44 |
| *Xgwm251* | 4B\4D | 152 | 184 | 0 | 3.05 |
| *Xgwm294* | 2A | 157 | 175 | 4 | 1.44 |
| *Xgwm297* | 7B | 174 | 162 | 0 | 0.43 |
| *Xgwm3* | 3D | 162 | 174 | 0 | 0.43 |
| *Xgwm304* | 5A | 175 | 155 | 6 | 2.01 |
| *Xgwm310* | NA | 171 | 165 | 0 | 0.11 |
| *Xgwm312* | 2A | 149 | 187 | 0 | 4.30 |
| *Xgwm333* | 7B | 175 | 158 | 3 | 1.19 |
| *Xgwm334* | 6A | 159 | 177 | 0 | 0.96 |
| *Xgwm337* | 1B\1D | 153 | 183 | 0 | 2.68 |
| *Xgwm350* | 4A | 188 | 148 | 0 | 4.76 |
| *Xgwm376* | 3B | 163 | 173 | 0 | 0.30 |
| *Xgwm389* | 3B | 163 | 173 | 0 | 0.30 |
| *Xgwm397* | 4A | 185 | 147 | 4 | 5.25 |
| *Xmag3810* | 7A | 173 | 163 | 0 | 0.30 |
| *Xpsp3000* | 1B\1D | 164 | 172 | 0 | 0.19 |
| *Xwmc1* | 6A | 163 | 173 | 0 | 0.30 |
| *Xwmc170* | 2A | 162 | 174 | 0 | 0.43 |
| *Xwmc177* | 2A | 184 | 152 | 0 | 3.05 |
| *Xwmc198* | 2A | 160 | 176 | 0 | 0.76 |
| *Xwmc222* | 1B\1D | 168 | 168 | 0 | 0.00 |
| *Xwmc238* | 4B | 156 | 180 | 0 | 1.71 |
| *Xwmc256* | 6A | 160 | 176 | 0 | 0.76 |
| *Xwmc283* | 7A | 188 | 148 | 0 | 4.76 |
| *Xwmc2830* | NA | 171 | 165 | 0 | 0.11 |
| *Xwmc296* | 2A | 166 | 170 | 0 | 0.05 |
| *Xwmc326* | 3B | 154 | 180 | 2 | 2.33 |
| *Xwmc335* | 7B | 172 | 164 | 0 | 0.19 |
| *Xwmc338* | 7B | 185 | 149 | 2 | 4.30 |
| *Xwmc364* | 7B | 174 | 162 | 0 | 0.43 |
| *Xwmc382* | 2A\2B | 181 | 152 | 3 | 3.05 |
| *Xwmc407* | 2A | 173 | 163 | 0 | 0.30 |
| *Xwmc415* | 5B | 155 | 181 | 0 | 2. 01 |
| *Xwmc44* | 1B | 153 | 183 | 0 | 2. 68 |
| *Xwmc47* | 4B | 175 | 161 | 0 | 0. 58 |
| *Xwmc476* | 7B | 173 | 163 | 0 | 0. 30 |
| *Xwmc487* | 6B | 160 | 176 | 0 | 0. 76 |
| *Xwmc494* | 6B | 161 | 175 | 0 | 0. 58 |
| *Xwmc503* | 2D | 184 | 152 | 0 | 3. 05 |
| *Xwmc506* | 7D | 180 | 153 | 3 | 2. 68 |
| *Xwmc532* | 3A | 161 | 175 | 0 | 0. 58 |
| *Xwmc533* | 3B\3D | 182 | 151 | 3 | 3. 44 |
| *Xwmc631* | 3D | 158 | 178 | 0 | 1. 19 |
| *Xwmc634* | 7D | 171 | 165 | 0 | 0. 11 |
| *Xwmc651* | 2A\3A | 167 | 169 | 0 | 0. 01 |
| *Xwmc696* | 7B | 170 | 166 | 0 | 0. 05 |
| *Xwmc710* | 4B | 150 | 186 | 0 | 3. 86 |
| *Xwmc720* | 4D | 172 | 164 | 0 | 0. 19 |
| *Xwmc728* | 1B | 170 | 166 | 0 | 0. 05 |
| *Xwmc752* | 5A | 185 | 146 | 5 | 5. 76 |
| *Xwmc758* | 7B | 177 | 159 | 0 | 0. 96 |
| *Xwmc765* | 5D | 175 | 161 | 0 | 0. 58 |
| *Xwmc766* | 1B | 180 | 156 | 0 | 1. 71 |
| *Xwmc773* | 6D | 172 | 164 | 0 | 0. 19 |
| *Xwmc783* | 5B | 163 | 173 | 0 | 0. 30 |
| *Xwmc93* | 1A\1D | 180 | 152 | 4 | 3. 05 |
| *Xgwm413* | 1B | 173 | 163 | 0 | 0. 30 |
| *Xgwm425* | 2A | 167 | 169 | 0 | 0. 01 |
| *Xgwm437* | 7D | 165 | 171 | 0 | 0. 11 |
| *Xgwm459* | 6A | 162 | 174 | 0 | 0. 43 |
| *Xgwm484* | 2D | 174 | 162 | 0 | 0. 43 |
| *Xgwm493* | 3B | 166 | 170 | 0 | 0. 05 |
| *Xgwm499* | 5B | 168 | 168 | 0 | 0. 00 |
| *Xgwm52* | 3D | 181 | 150 | 5 | 3. 86 |
| *Xgwm566* | 3B | 173 | 163 | 0 | 0. 30 |
| *Xgwm583* | 5D | 168 | 168 | 0 | 0. 00 |
| *Xgwm617* | 5A\6A | 153 | 183 | 0 | 2. 68 |
| *Xwmc710* | 4B | 151 | 185 | 0 | 3. 44 |

We carried out further research on 1AS. We first developed 13 pairs of new SSR markers located on 1AS (Table 2). Of these, nine pairs of markers (*SSR11, SSR26, SSR110, SSR113, SSR114, SSR115, SSR122, SSR144*, and *SSR365*) could be used to detect the sequence elimination. SSR markers are co-dominant, and polyacrylamide gel electrophoresis (PAGE) can be used to distinguish different genotypes due to molecular weight differences. In this study, SSR markers were able to detect the two parental genotypes and sequence elimination. The results with *SSR11* are shown in Figure 1A as an example. A 1AS genetic linkage map was constructed using the RIL population (Figure 2). The genetic linkage map indicated that the SSR markers that could detect the sequence elimination are concentrated at the end of 1AS. We also found that the junction of the sequence elimination chromosome and the normal chromosome is located between *SSR110* and *SSR283;* the genetic distance between these two markers is 0.98 cM.

**Figure 1.**
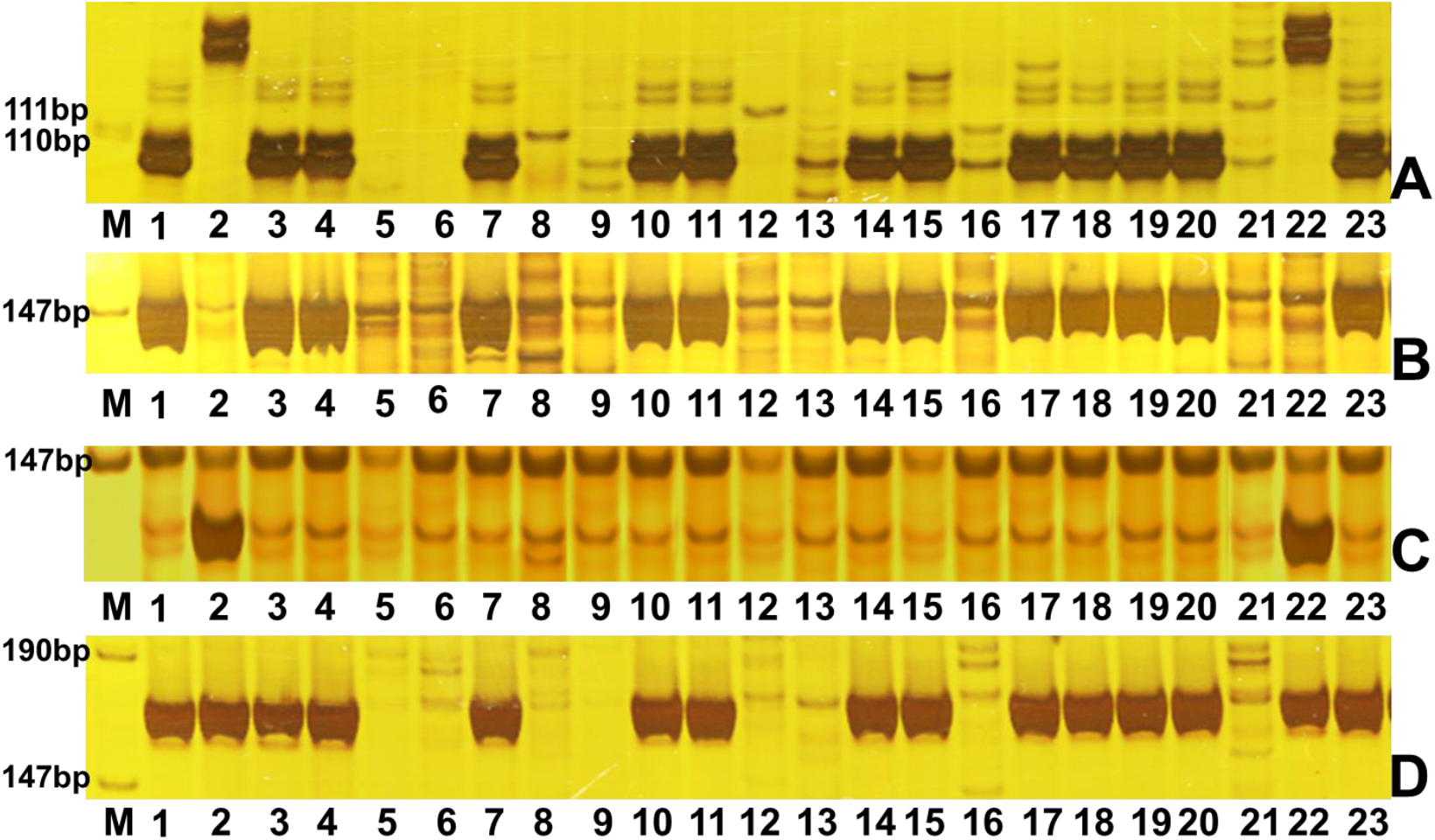
Detection of different markers in parents and RILs. A, PAGE result of the co-dominant SSR marker *SSR11*. B, PAGE result of the Pubing3504 dominant SNP marker *SNP5*. C, PAGE result of the Jing4839 dominant SNP marker *SNP_174*. D, PAGE result of primer *SNP_SE4*. M is marker lane; lane 1 is Pubing3504; lane 2 is Jing4839. Lanes 3 to are RILs. Lanes 5, 6, 8, 9, 12, 13, 16, and 21 are sequence elimination lines.

**Figure 2.**
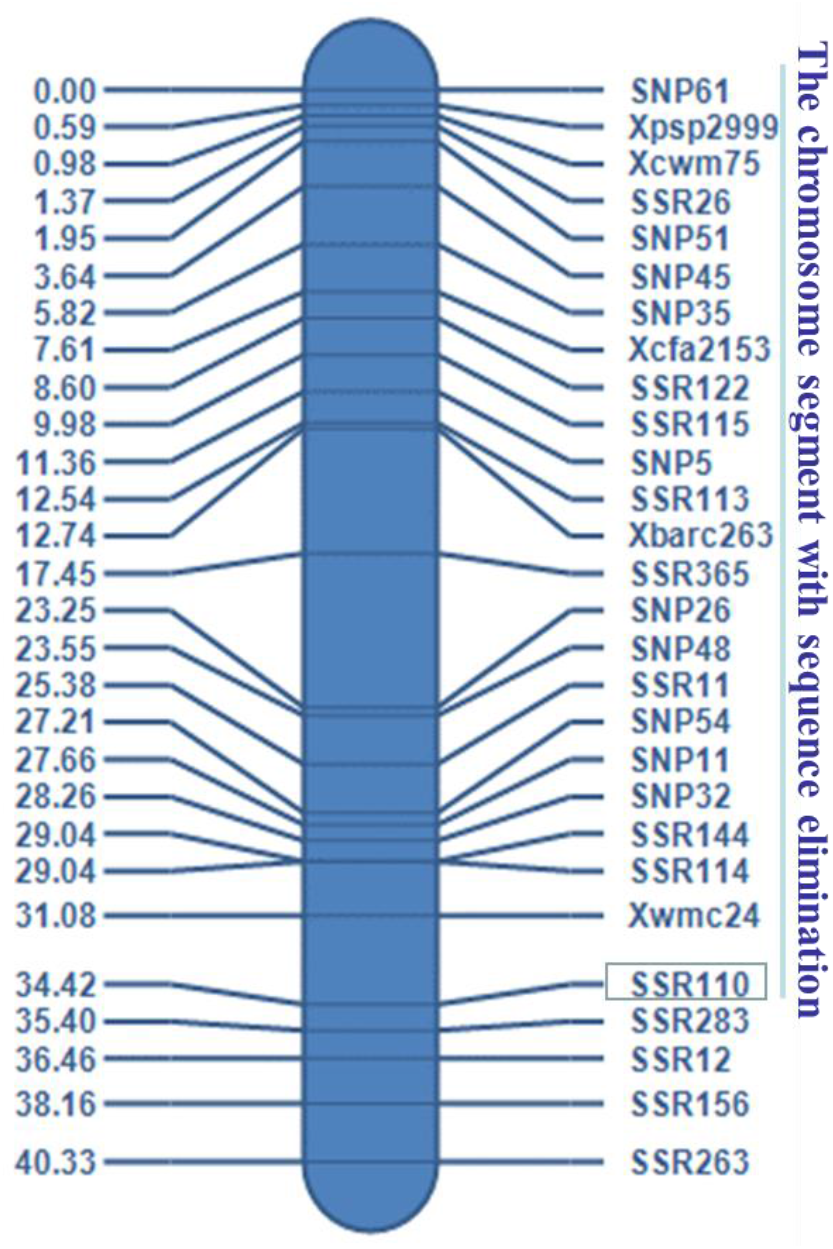
Genetic linkage map of 1AS constructed using the RIL population. Sequence elimination occurred in the chromosome segment from *SSR110* (boxed) to the end of 1AS. This chromosome segment includes 14 pairs of co-dominant SSR markers and 10 pairs of dominant SNP markers of Pubing3504.

**Table 2.**
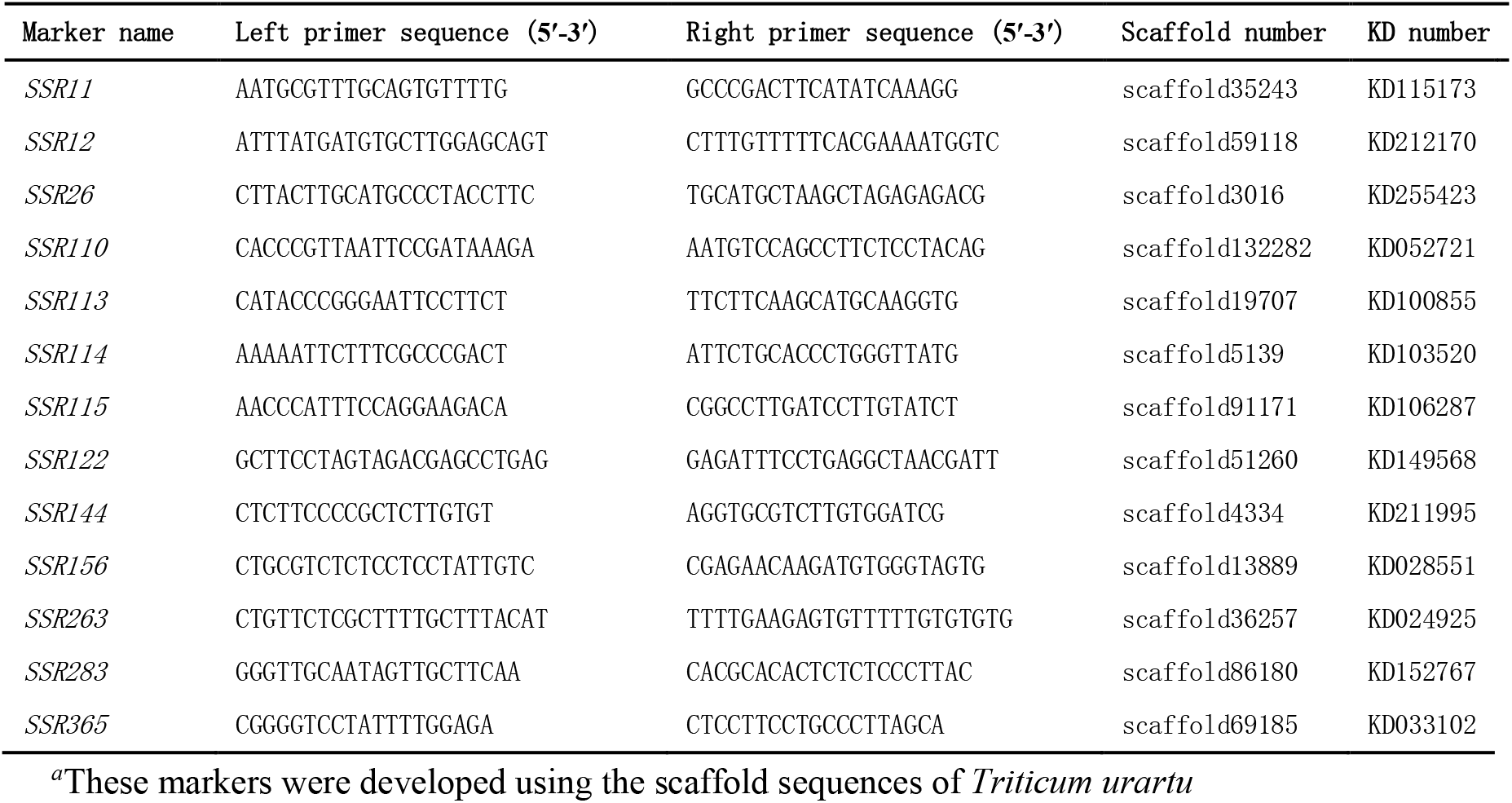
The co-dominant SSR markers in 1AS developed in this research^*α*^.

### The origin of the chromosome segment with sequence elimination

We carried out further studies to determine the origin of the chromosome segment with sequence elimination. Theoretically, it may originate from Pubing3504, from Jing4839, or both. First, we developed dominant SNP markers for Pubing3504 using AS-PCR technology. Ten pairs of SNP markers (*SNP5, SNP11, SNP26, SNP32, SNP35, SNP45, SNP48, SNP51, SNP54*, and *SNP61*) were assessed and added to the chromosome segment of sequence elimination in the genetic linkage map (Figure 2, Table 3). Sequence elimination did not affect the genetic mapping of these SNP markers. The Jing4839 genotype and sequence elimination both presented as a missing genotype when using these SNP markers to detect RILs. The results with *SNP5* are shown in Figure 1B as an example.

**Table 3.**
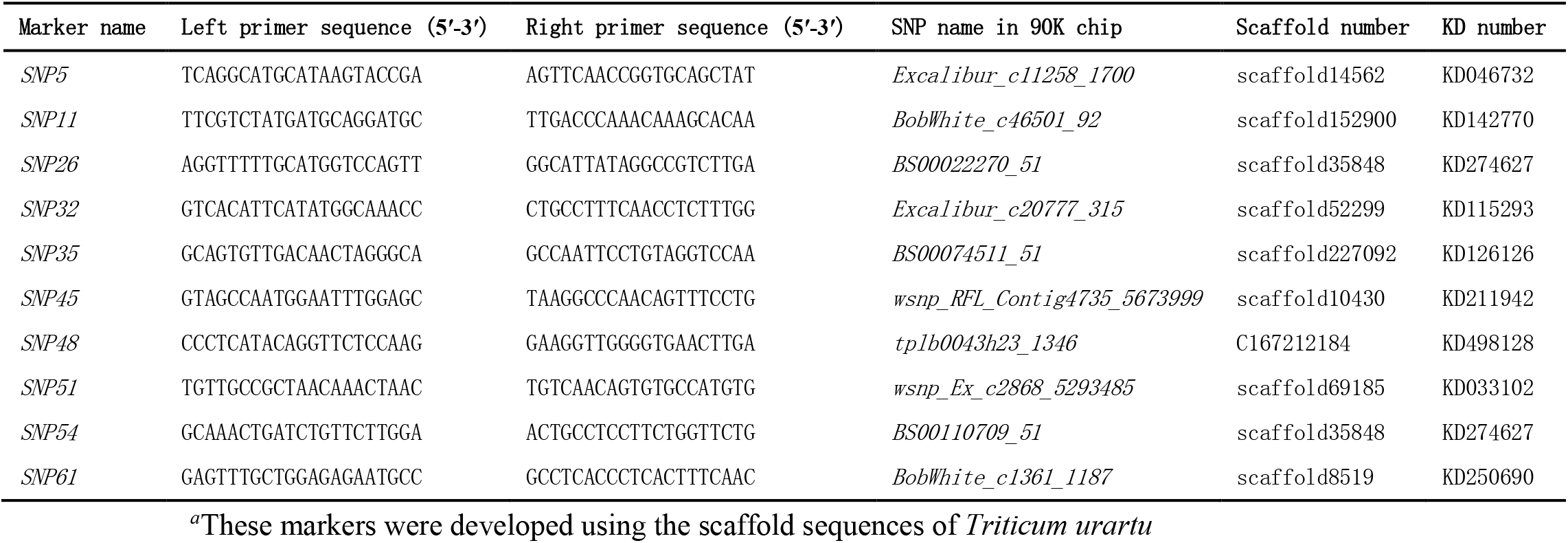
The dominant SNP markers of Pubing3504 in 1AS developed in this research^*α*^.

We then developed ten pairs of dominant SNP markers for Jing4839 using AS-PCR technology. They are *SNP_174, SNP_035, SNP_054, SNP_009, SNP_095, SNP_097, SNP_102, SNP_120, SNP_069*, and *SNP_024* (Table 4). These SNP loci were derived from the coding sequences on 1AS. The Pubing3504 genotype and sequence elimination both presented as a missing genotype when using these SNP markers to assess the RIL population. The results with *SNP_174* are shown in Figure 1C as an example. However, because so many of the RILs presented as missing genotype, these markers could not be mapped to the genetic linkage map. Therefore, it is reasonable to consider Jing4839 and sequence elimination as the same genotype to construct the genetic linkage map. Judging from these results, the chromosome segment with sequence elimination originated from Jing4839.

**Table 4.**
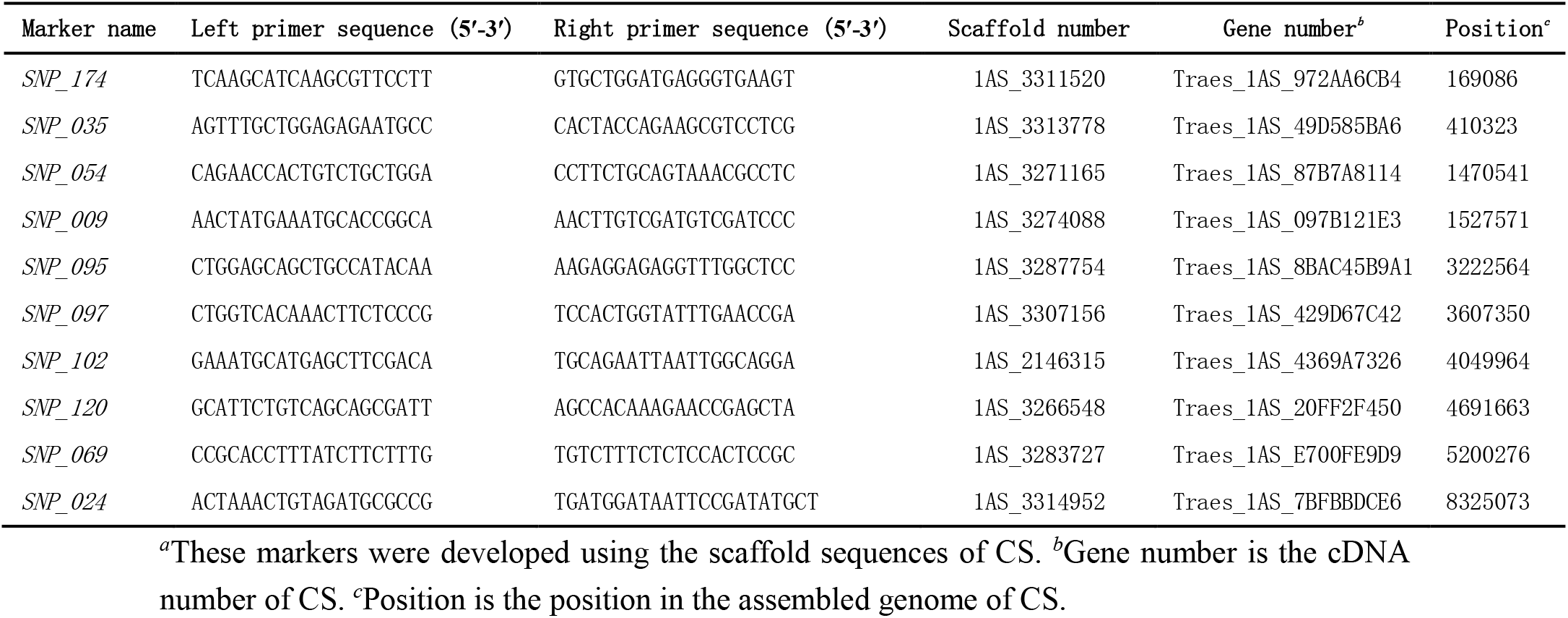
The dominant SNP markers of Jing4839 in 1AS developed in this research^*α*^.

### Region selectivity of sequence elimination

This study found that the sequence elimination frequently occurred at an SSR locus. To study the sequence elimination of SNP loci, we designed ten pairs of primers to amplify the sequence flanking these SNP sites (Table 5). Three pairs of these primers (*SNP_SE1, SNP_SE4*, and *SNP_SE7*) could be used to detect sequence elimination. These primers effectively amplified genomic DNA of the parental and offspring lines with no sequence elimination, and offspring lines with sequence elimination were indicated as genotype missing. The results with *SNP_SE4* are shown in Figure 1D as an example. These diagnostic primers indicated that sequence elimination occurred in 76 lines of the RIL population. These 76 lines were used in the following studies.

**Table 5.**
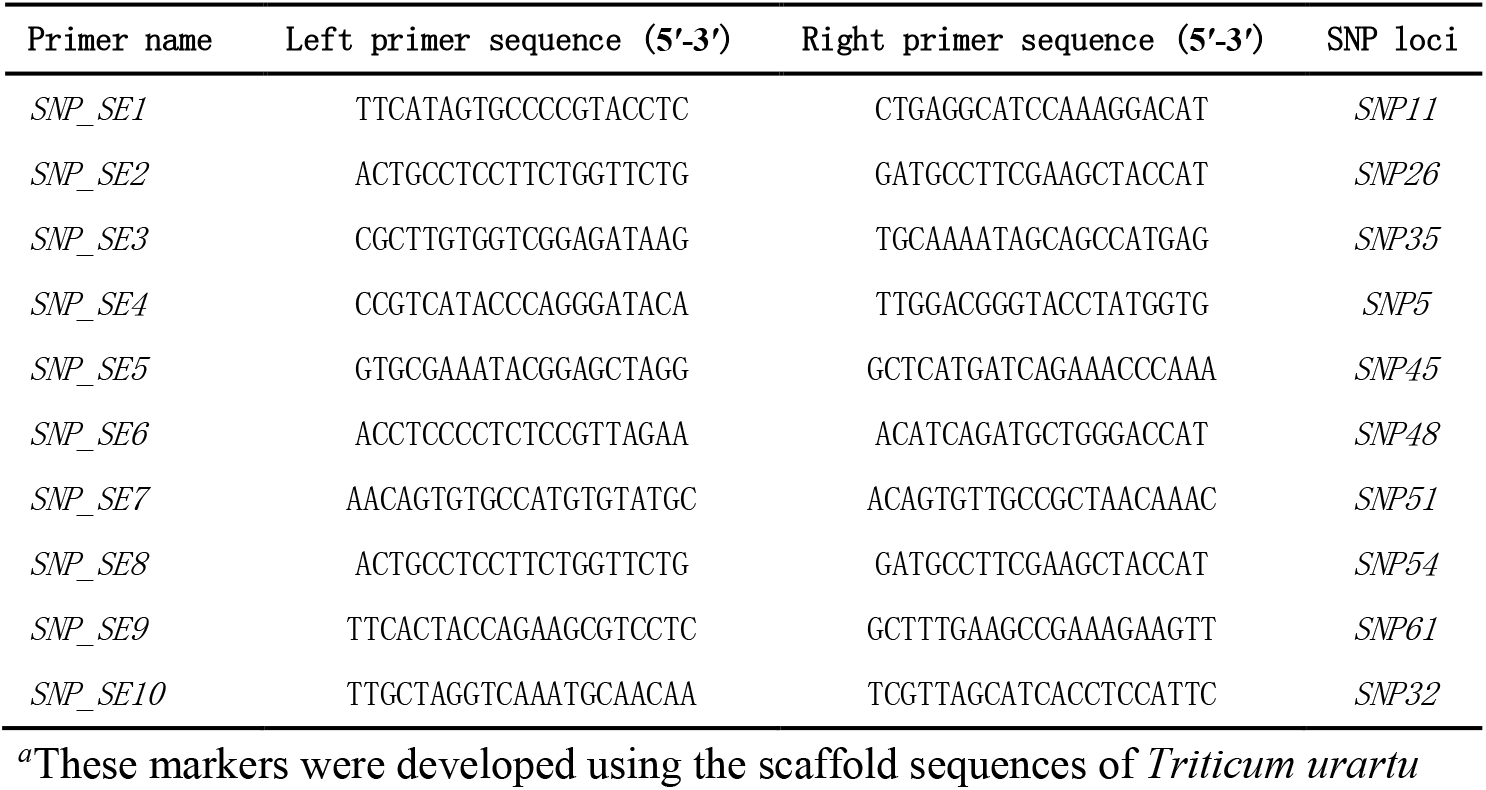
Primers used to detect sequence elimination of SNP loci^*α*^.

We further studied the sequence elimination on gene loci. RNA-sequencing was used in the two parents and a sequence elimination line from RILs. The results showed that some genes in 1AS were expressed in both parental transcriptomes, but were not expressed in the sequence elimination line. Ten gene loci with sequence elimination are shown in Table 4. These gene sequences have SNPs in common between the parents. We developed Jing4839 dominant SNP markers using these gene sequences. These markers were used to detect gene loci in the 76 RIL lines with sequence elimination, and we found that some gene sequences were eliminated from 1AS. Thus, sequence elimination can affect gene coding sequences in the genome and hence may alter gene expression.

### Genetic characteristics of the chromosome segment with sequence elimination

We examined the reorganization in the 336 lines of the RIL population using markers in the chromosome segment with sequence elimination. This chromosome segment in the genetic linkage map contains 14 pairs of SSR markers and 10 pairs of SNP markers (Figure 2). Recombination in the chromosome segment occurred in 60 lines, and recombination did not occur in 276 lines. We found that 184 of the 276 chromosome segments were from Pubing3504, 16 were from Jing4839, and 76 were from sequence elimination. Recombination only occurred between the homologous chromosomes of Pubing3504 and Jing4839 in the 60 RILs.

Finally, we established a genetic model to illustrate the sequence elimination that occurred in hybrid offspring of Pubing3504 and Jing4839 (Figure 3). F1 was obtained by Pubing3504 and Jing4839 hybridization. After F1 selfing, a new chromosome segment with sequence elimination was generated. The three chromosome segments could form six possible chromosome combinations. The combination of Pubing3504 and Jing4839 could undergo normal homologous recombination; however, the chromosome with sequence elimination could not undergo homologous recombination with either of the parentally derived chromosome segments.

**Figure 3.**
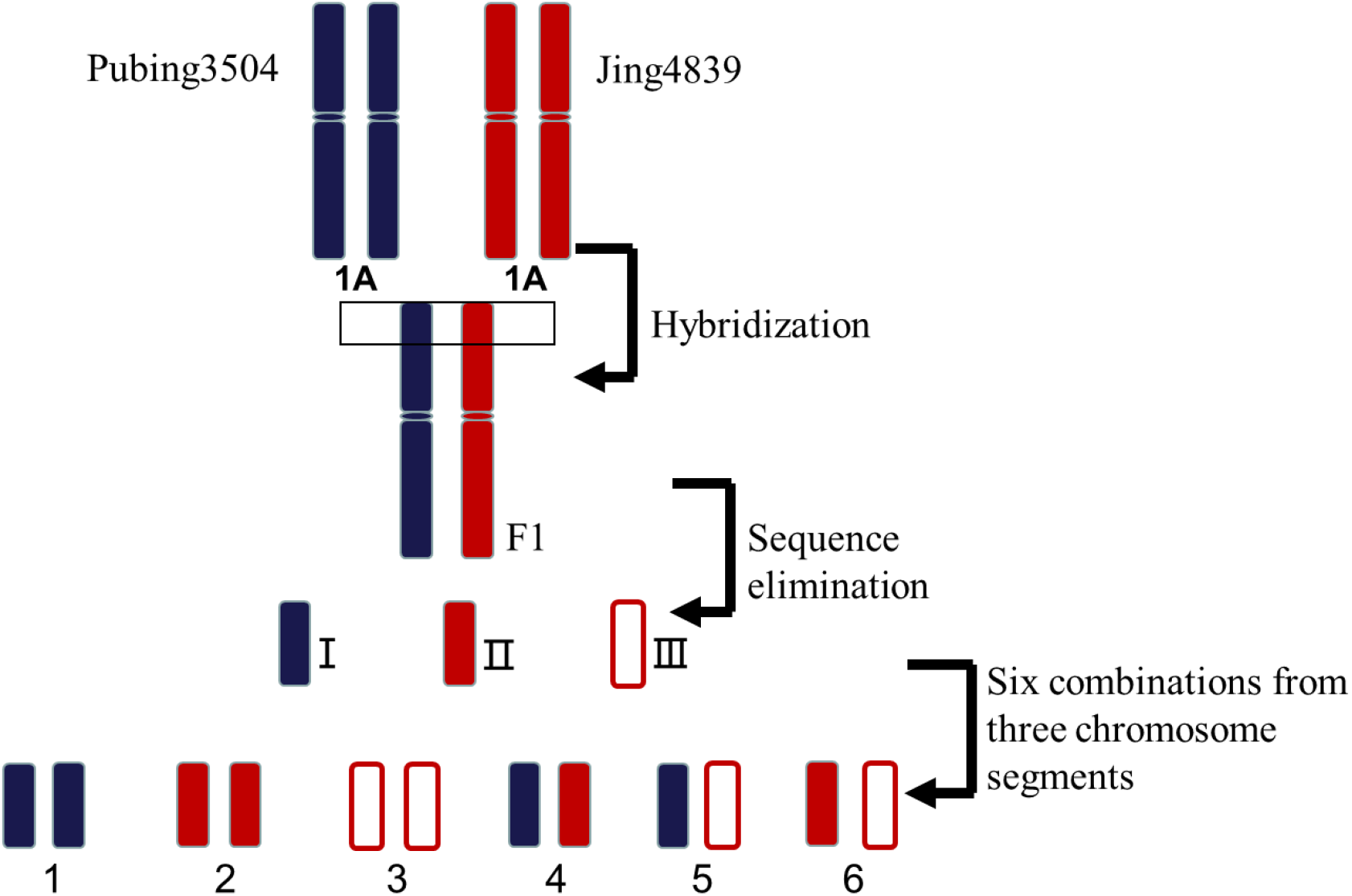
Genetic model for the sequence elimination examined in this research. F1 was obtained by Pubing3504 and Jing4839 hybridization. After F1 selfing, a new chromosome segment with sequence elimination was generated. The chromosome segments from Pubing3504 (blue) and Jing4839 (red) and the new chromosome segment with sequence elimination (white) are boxed. These chromosome segments could form six combinations; combinations 1–3 are homozygous, combinations 4–6 are heterozygous. Combination 4 (between Pubing3504 and Jing4839) can undergo normal homologous recombination, but combination 5 (between Pubing3504 and sequence elimination) and combination 6 (between Jing4839 and sequence elimination) cannot.

## DISCUSSION

### Characteristics of sequence elimination

Allopolyploidization causes rapid changes in the wheat genome, which improves the speed of evolution. These rapid changes include non-random sequence elimination in coding and non-coding DNA sequences, epigenetic changes in coding and non-coding DNA sequences, and changes in the activity of transposable elements affecting the expression of adjacent genes (Feldman and Levy 2005; Feldman and Levy 2009; Feldman and Levy 2012).

Sequence elimination is a major genomic change resulting from allopolyploidization. Eliminated sequences include both repeat and low-copy DNA sequences (Han et al. 2005; Guo and Han 2014). In the wheat genome, more than 80% of the DNA sequence is repeat sequence (Gustafson et al. 2007; Wicker et al. 2011). Because of this, sequence elimination can cause a reduction in the amount of genomic DNA. Some studies have shown that the amount of genomic DNA in common wheat and synthetic hexaploid wheat is less than the total amount of the parents (Ozkan et al. 2003; Gustafson et al. 2008; Eilam et al. 2010). Low-copy sequences in the wheat genome can be divided into genome-specific sequences found only in the A, B, or D genome and chromosome-specific sequences, for example, only in the 1A, 1B, and 1D chromosomes. Some studies found that some chromosome-specific sequences are eliminated from a genome or two genomes in common wheat and synthetic hexaploid wheat compared with their diploid progenitors (Feldman et al. 1997; Liu et al. 1998b; Ozkan et al. 2001).

In this study, we found that sequence elimination occurred in the hybrid offspring of Pubing3504 and Jing4839. We observed sequence elimination, all occurring at the end of 1AS. The eliminated sequences we found are low-copy sequences in the genome. These sequences are both A genome–specific sequences and 1A chromosome-specific sequences, which are commonly used for marker development to construct the genetic linkage map. We observed sequence elimination in both SSR loci and SNP loci. At these loci, there are base differences between Pubing3504 and Jing4839. Thus, base differences between the parents may trigger the sequence elimination. We also found that sequence elimination occurred in the gene coding sequence. Some gene coding sequences were eliminated from the genome through sequence elimination. These genes could be expressed in the transcriptome of Pubing3504 and Jing4839, but were not expressed in the transcriptome of the line with sequence elimination. Therefore, sequence elimination can affect gene expression.

Sequence elimination is asymmetric or preferential to different genomes. In the octoploid triticale genome (AABBDDRR), the wheat genome (AABBDD) is conservative, and the rye genome (RR) has undergone great changes (Ma et al. 2004). Some studies have indicated that sequence elimination tends to be in the larger genome (Schwarzacher et al. 2011). The preference of sequence elimination results in gene expression as a diploid pattern (Le Comber et al. 2010; Qi et al. 2012). In this study, we found that the sequence elimination was also asymmetric.

Sequence elimination usually occurs in the early generations after hybridization. Previous studies showed that the sequence elimination may take place in the F1 generation after hybridization (Salina et al. 2004). We previously demonstrated sequence elimination in F2 after hybridization between Pubing3504 and Jing4839 (Chen 2012). In the F1 generation, the chromosome doubling is not yet complete (Ozkan et al. 2001; Han et al. 2003; Ma and Gustafson 2006). Therefore, the elimination of sequences is not caused by chromosome doubling. Some studies suggest that sequence elimination may be caused by DNA methylation (Han et al. 2003; Guo and Han 2014). The reason for the sequence elimination is not yet definitively known.

### Sequence elimination causes chromosome differentiation

The genome of common wheat is composed of seven parts of homologous groups. For example, the first homologous group includes the 1A, 1B, and 1D chromosomes. All chromosomes in a homologous group have a common ancestor, so they have similar gene order and DNA sequences. Although they are homologous chromosomes, they do not pair during meiosis. One reason for this is that a major gene *Ph1*, which is mapped in 5BL, allows homologous chromosome pairing from the same genome (for example between 1A and 1A) and inhibits non-homologous chromosome pairing and homologous chromosome pairing from different genomes (for example between 1A and 1B, between 1A and 1D, or between 1B and 1D) that have evolved in the common wheat genome. So far, the mechanism by which *Ph1* controls homologous chromosome pairing is not clear (Sears 1976).

Sequence elimination is another mechanism to inhibit homologous chromosome pairing from different genomes in common wheat, which causes differentiation of homologous chromosomes from different genomes to inhibit chromosome pairing between them (Feldman et al. 1997; Ozkan et al. 2001; Feldman and Levy 2012). In previous studies, 18 newly formed allopolyploids were used to study chromosome pairing in meiosis and seed fertility in S1, S2, and S3 generations; the results show that the number of bivalents in meiosis and the seed fertility increase with the increase of the generation of the allopolyploid. The cytological behavior of the allopolyploid was considered to begin as the diploid in early generations after chromosome differentiation caused by sequence elimination (Gustafson et al. 2009). In our research, we used high-density molecular markers to detect the chromosome segment with sequence elimination in RILs, and we found that there was no homologous recombination in this chromosome segment. We think that the sequence elimination caused the differentiation of chromosomes, and the chromosome differentiation affected the homologous pairing at this chromosome segment in meiosis, which further affected the occurrence of homologous recombination at this chromosome segment.

## Acknowledgments

This work was supported by the Agricultural Science and Technology Innovation Program of CAAS and the National Key Technologies R&D Program of China during the 12th Five-Year Plan period (grant No. 2013BAD01B02).

